# Climate as a driver of adaptive variations in ecological strategies in *Arabidopsis thaliana*

**DOI:** 10.1101/404210

**Authors:** François Vasseur, Kevin Sartori, Etienne Baron, Florian Fort, Elena Kazakou, Jules Segrestin, Eric Garnier, Denis Vile, Cyrille Violle

## Abstract

**Background and aims:** The ‘CSR classification’ categorizes plant species between stress-tolerators, ruderals (R) and competitors (C). Initially proposed as a general framework to describe ecological strategies at the interspecific level, this scheme has recently been used to investigate the variation of strategies within species. For instance, ample variation along the S-R axis was found in *Arabidopsis thaliana*, with stress-tolerator accessions predominating in hot and dry regions.

**Methods:** In this study, the range of CSR strategies within *A. thaliana* was evaluated across 426 accessions originating from North Africa to Scandinavia. A position in the CSR strategy space was allocated for every accession based on three functional traits: leaf area, leaf dry matter content (LDMC) and specific leaf area (SLA). Results were related to climate at origin and compared to a previous study performed on the same species. Furthermore, the role of natural selection in phenotypic differentiation between lineages was investigated with Q_st_-F_st_ comparisons, using the large genetic information available for this species.

**Key results:** Substantial variation in ecological strategies along the S-R axis was found in *A. thaliana*. By contrast with previous findings, stress-tolerator accessions predominated in cold climates, notably Scandinavia, where late flowering was associated with traits related to resource conservation such as high LDMC and low SLA. Because of trait plasticity, variations in CSR classification to growth conditions were also observed for the same genotypes.

**Conclusions:** There is a latitudinal gradient of ecological strategies in *A. thaliana* as a result of within-species adaptation to climate. Our study also underlines the importance of growth conditions and of the methodology used for trait measurement, notably age versus stage measurement, to infer the strength and direction of trait-environment relationships. Taken together, this highlights the potential and limitations of the CSR classification to explain functional adaptation to the environment.

## Introduction

Screening approaches allow species comparison on the basis of key ‘functional traits’, *i.e*. traits representative of major functions such as growth, stress resistance, defence and reproduction (Keddy 1992, Violle *et al.* 2007). Trait-based approaches in plant ecology have a long history of classifying plant species into functional groups according to the combination of phenotypic traits they exhibit (Garnier *et al.* 2016). Such approaches have been mainly applied for comparative analyses at the interspecific level to identify general patterns of trait variation and covariation. However, recent comparative analyses argue for a better integration of intraspecific variability for understanding the role of trait covariation in plant adaptation, ecosystem functioning and community assembly (Albert *et al.* 2010, 2011; Violle *et al.* 2012; Siefert *et al.* 2015).

Amongst the prominent examples of plant species classification, Grime (1977) defined ecological strategies based on the idea that there are two main ecological drivers of plant diversification: (i) the effect of stress related to the shortage of resources (*e.g.*, nutrient, water and light), and (ii) the effect of disturbance. Stress is viewed in this context as any environmental factors or combination of factors that reduce plant growth, although the shortage of nutrients, water or light can each affect specific traits (Grime and Hunt, 1975; Grime 1977; Hodgson *et al*. 1999). By contrast, disturbance is viewed as factors that cause the partial or total destruction of plant biomass, which include grazing, trampling, mowing, but also extreme climatic events such as severe drought, frost and fire (Grime and Hunt, 1975). Differences in disturbance and stress intensity are expected to result in quantitative variation in three ecological strategies: (i) stress-tolerators (S) in stressed, resource-poor habitats with low disturbance, which invest resources to protect tissue from stress damages, (ii) ruderals (R) in resource-rich environments associated with repeated disturbance, which invest resources into rapid reproduction and propagule dispersal, and (iii) competitors (C) in highly productive habitats with low stress intensity and disturbance, which invest resources into the rapid growth of large organs to outcompete neighbours. The S-R axis is traditionally viewed as an axis of resource-use variations at the leaf level (Pierce *et al.* 2013), where ruderality is associated with acquisitive resource-use (characterized by short-lived, flimsy leaves with high nutrient concentration and high net photosynthetic rate), and stress-tolerance is associated with conservative resource-use (characterized by long-lived, tough leaves with low nutrient concentration and low net photosynthetic rate). By contrast, variation of competitive ability along the C axis is thought to reflect variation in plant and organ size, and it is expected to operate where the impacts of stress and disturbance are low (Grime 1977; Hodgson *et al*. 1999).

Originally designed in the context of temperate herbaceous vegetation, the CSR scheme has been extended to other types of vegetation (Caccianiga *et al.* 2006; Navas *et al.* 2010; Schmidtlein *et al.* 2012), including a recent worldwide application (Pierce *et al.* 2017). An algorithm has recently been developed to quantify the CSR scores of diverse plant species based on the measurement of three leaf traits: leaf area (LA), specific leaf area (SLA) and leaf dry matter content (LDMC) (Pierce *et al.* 2013, 2017). Albeit less precise than methods that consider whole-plant traits which are more closely associated with stress response, competitive ability and ruderality (Hodgson *et al.* 1999), classification tools based on few leaf traits have the advantage that many measurements can be performed with minimal effort. This allows comparing very ecologically disparate species (Pierce *et al.* 2017), or many genotypes and populations within species (May *et al.* 2017).

*Arabidopsis thaliana* is a small rosette-shaped species that is widely used in molecular biology and quantitative genetics. It has recently gained a renewed interest in evolutionary ecology due to the large collection of natural accessions collected from various climates and genotyped at high density (Weigel 2012). Furthermore, *A. thaliana* has been shown to exhibit a significant range of phenotypic variation in relation to climate, making it possible to investigate the genetic and evolutionary drivers of functional diversification (Vasseur *et al*. 2018). For instance, Q_st_-F_st_ analysis has been proposed as a powerful way to discriminate adaptive and non-adaptive processes at the origin of phenotypic differentiation between genetic groups, populations or lineages (Leinonen *et al*. 2013). Indeed, this method allows one to compare the level of phenotypic differentiation (Q_st_) to the genetic differentiation (F_st_) expected under the neutral model of population divergence. In plants, this has been used to investigate the role of selection at the origin of between-populations phenotypic differences related to resource-use traits (Brouillette *et al.* 2014), drought resistance (Ramírez-Valiente *et al.* 2018), life history traits (Moyers and Rieseberg 2016), and functional adaptation to an elevation gradient (Luo *et al.* 2015).

*A.thaliana* is generally described as a ruderal species that, like most annual plants, reproduces quickly and invests preferentially resources to the production and dispersal of propagules (Díaz *et al.* 2016; Pierce *et al.* 2017). In a recent paper, May *et al*. (2017) used the CSR framework to investigate intraspecific variation in ecological strategies within this species. Using 16 accessions originating from contrasted climates in Europe, they found that *A. thaliana* actually exhibits a wide range of variation from ruderals to stress-tolerators, with most accessions being classified as intermediate (SR) and none as competitor. Interestingly, May *et al.* also found that ruderality was negatively correlated with the temperature at the site where the accession originated from. For instance, stress-tolerators originated predominantly from sites in hot climates (Libya, Sicily and Cape Verde Islands). However, May *et al.* (2017) used a relatively low number of accessions, which prevents examining the evolutionary and adaptive bases of CSR variations with the environment.

In the present study, we analysed CSR variations on a set of 426 *A. thaliana* accessions originating from contrasting climates in Europe, North Africa and East Asia. Using the classification method based on three leaf traits (area, SLA and LDMC) (Pierce *et al.* 2017), we tested the range of ecological strategies exhibited by these accessions. We investigated whether the variation in strategies can be attributed to adaptive processes, using the genetic data available in this species to perform Q_st_-F_st_ analysis. We also examined how CSR strategies measured with leaf traits correlated with whole-plant traits related to competitive ability (rosette size) and propagule dispersal (fruit number). Finally, we compared our results to the findings from May *et al.* (2017), and we discussed the possible causes of differences between studies such as the direction of trait-environment relationships.

## Materials and Methods

### Plant material

Two experiments were performed in this study: the first one in the PHENOPSIS automaton (see below) and the second one in greenhouse. In the first experiment, we used a total of 400 natural accessions of *A. thaliana* L. Heynh representative of a geographical sampling from the worldwide lines of the RegMap population (Horton *et al.* 2012) (*n* = 214) and from French local populations (Brachi *et al.* 2013) (*n* =186). In the second experiment, we used a total of 200 accessions from a random sampling from the worldwide lines of the RegMap population. Overall, 426 accessions ranging latitudinally from North Africa to Scandinavia were phenotypically characterized, 172 of which were common to both experiments (Tables S1 and S2).

### Experimental design

In Experiment 1 (PHENOPSIS), plants were grown in the high-throughput phenotyping platform PHENOPSIS (Granier *et al.* 2006) in 2014, using one replicate plant per accession, except for Col-0 for which there were 10 replicates. Seeds were stratified in the dark at 4 °C for at least one week before sowing to ensure homogeneous germination among genotypes. Four to six seeds were sown at the soil surface in 225 ml pots filled with a 1:1 (v:v) mixture of loamy soil and organic compost. Prior to sowing, soil surface was moistened with one-tenth strength Hoagland solution, and pots were kept in the dark for 48 h under controlled environmental conditions (20 °C, 70% air relative humidity). Pots were then placed in the PHENOPSIS automaton growth chamber at 20 °C, 12 h day length, 70 % relative humidity, 175 μmol m^−2^ s^−1^ PPFD. Pots were sprayed with deionized water three times per day until germination, soil water content was then adjusted to 0.35 g H_2_O g^−1^ dry soil (–0.07 MPa soil water potential) to ensure optimal growth (Aguirrezábal *et al.* 2006; Vile *et al.* 2012; Vasseur *et al.* 2014). After emergence of the fourth leaf, seedlings were thinned to keep only one plant in each pot.

In Experiment 2 (Greenhouse), plants were grown in four replicates per accession in greenhouse between December 2015 and May 2016. Seeds were sown on organic soil and stratified at 4 °C for four days. At the emergence of the first two true leaves, plants were transplanted in 300 ml individual pots filled with a 1:1 (v:v) mixture of loamy soil and organic compost. Pots were randomly distributed among four blocks that were rotated every day in the greenhouse. All pots were watered twice a week. To reduce environmental heterogeneity in the greenhouse, walls were painted in white and a semi-transparent curtain was installed below the glass roof. Additional light was provided to reach *ca*. 65 μmol m^−2^ s^−1^ PPFD. Photoperiod and temperature were kept constant at 12 h day length, and 18/16 °C day/night, respectively.

### Trait measurement

In both experiments, traits were measured following standardized protocols (Perez-Harguindeguy *et al.* 2013) at a fixed phenological stage when flower buds were macroscopically visible (*i.e*. bolting stage, used as measurement of flowering time, FT). The lamina of a fully-expanded, adult and non-senescent leaf exposed to light was detached from the rosette, kept in deionised water at 4 °C for 24 h for water saturation, and then weighted (mg). After the determination of water-saturated mass, individual leaves were scanned for determination of leaf lamina area (LA, mm^2^) using ImageJ (https://imagej.nih.gov/ij/). Dry mass of the leaf lamina was obtained after drying for 72 h at 65 °C. Leaf dry matter content (LDMC, mg g^−1^) and specific leaf area (SLA, mm^2^ mg^−1^) were calculated as the ratio of lamina dry and water-saturated mass, and the ratio of lamina area to lamina dry mass, respectively (Perez-Harguindeguy *et al.* 2013). In PHENOPSIS, plants were harvested at first opened flower and rosette fresh mass (mg) was measured. In Greenhouse, plants were harvested after full senescence and total number of fruits was manually counted on the inflorescence. Overall, out of the 400 and 200 accessions in PHENOPSIS and Greenhouse, respectively 357 and 198 accessions were completely phenotyped for all traits (Tables S1 and S2), with 152 accessions common to both experiments.

We calculated CSR scores (*i.e.* % along C, S and R axes) for all accessions in PHENOPSIS and Greenhouse based on three traits: LA, LDMC and SLA, using the recent method developed by Pierce *et al.* (2017). The method is based on an algorithm which combines data for three leaf traits (LA, SLA and LDMC) that were shown to reliably position the position of species in the CSR scheme. We calculated CSR scores for each accession using average trait value per experiment using the calculation table provided in Supplementary Information of Pierce *et al.* (2017).

### Re-analysis of published data

In our study, there were several accessions common with a previously published analysis of CSR variations in *A. thaliana* (May *et al*. 2017). Ten accessions were common between May *et al*. and the PHENOPSIS experiment on the one hand, and six accessions with the greenhouse experiment on the other hand. In May *et al*. (2017), CSR scores were calculated based on six traits with a method previously proposed by Hodgson *et al*. (1999). To compare both datasets, we first recalculated CSR scores from data in May *et al*. with Pierce’s method, using LA, LDMC and SLA provided for their 16 accessions (May *et al*. 2017), and compared them to the CSR scores they measured with the Hodgson’s method.

### Genetic analysis and Q_st_-F_st_ comparisons

Genetic groups in *A. thaliana* were determined by clustering of 395 accessions for PHENOPSIS dataset, and 198 accessions for Greenhouse dataset, both using the 250K Single Nucleotide Polymorphisms (SNPs) data available from Horton *et al*. (2012). Clustering was performed with ADMIXTURE (Alexander *et al.* 2009) after linkage disequilibrium pruning (*r*^2^ < 0.1 in a 50 kb window with a step size of 50 SNPs) with PLINK (Purcell *et al.* 2007), resulting in 24,562 independent SNPs. We assigned each genotype to a group if more than 60% of its genome derived from the corresponding cluster. The accessions not matching this criterion were labelled “Admixed” and were not used for the F_st_ and Q_st_ calculation. Cross-validation for different number of genetic clusters revealed that the PHENOPSIS dataset was composed of six genetic groups (group 1 = 74 accessions, group 2 = 48, group 3 = 18, group 4 = 55, group 5 = 5, group 6 = 71, admixed = 123), while the Greenhouse dataset was composed of four genetic groups (group 1 = 38 accessions, group 2 = 16, group 3 = 83, group 4 = 7, admixed = 54). Consistent with the hypothesis of genetic divergence because of isolation by distance, these genetic groups were geographically clustered (Fig. S1). We calculated Weir and Cockerham F_st_ value for all the 24,562 SNPs, and Q_st_ as the between-group phenotypic variance divided by the total phenotypic variance, using mixed-effect models with Group as random factor. We used parametric bootstrap method to generate 95% confidence intervals (CI) around Q_st_ values with the package ‘*MCMCglmm*’ in R (100,000 iterations).

### Statistical analyses

Genotypic means in the greenhouse experiment were estimated as the fitted genotypic values from the linear models, using *lsmeans* function. Genotype effect on trait variation and broad-sense heritability (*H*^2^) were assessed using individual data from the greenhouse experiment (Table S3). Genotype effect was tested with one-way ANOVA following linear modelling, using Genotype and Block as explanatory variables. *H*^2^ was measured as the ratio of phenotypic variance attributable to genotypic effect over total phenotypic variance, using mixed-effect models with Block as fixed factor and Genotype as random factor, using the package *nlme* in R.

Climate variables at the collection points of each accession were extracted from the Worldclim database (http://www.worldclim.org/bioclim), with a 2.5 arc-minutes resolution. Trait-trait, trait-environment and trait-CSR relationships were examined with Spearman’s rank coefficients of correlation (ρ), and associated *P*-values (*P*), using the function *cor.test* (Table S4). Pearson’s coefficients of correlations (*r*) between traits and climatic variables were also calculated (Table S5). Regression lines were drawn from Standard Major Axis (SMA), using the package *smatr*. All analyses were performed in R 3.2.3 (Team RC 2014).

## Results

### Trait variation and covariation

All traits varied significantly among accessions (all *P* < 0.001, Table S3). We found that FT ranged between 30 and 101 days (57 days on average) in PHENOPSIS, and between 25 to 115 days (61 days on average) in Greenhouse. Trait variation was mainly due to genetic variability among accessions, as measured by the high amount of phenotypic variance accounted for genotype effect (*H*^2^ ranged between 0.58 for LA and 0.73 for SLA, 0.88 for FT; Table S3). Most traits were correlated with each other (Fig. S2; Tables S4, S5): SLA and LDMC were negatively correlated (Spearman’s coefficient ρ = −0.94 and −0.88 in PHENOPSIS and Greenhouse, respectively, both *P* < 0.001; Fig. S2F), and FT was positively correlated with LDMC (ρ = 0.63 and 0.86, *P* < 0.001; Fig. S2B), and negatively with SLA (ρ = −0.73 and −0.92, both *P* < 0.001; Fig. S2D).

### CSR classification

*A. thaliana* accessions mainly varied along the S-R axis, between purely ruderals (R) to moderate stress-tolerators (S/SR) (Fig. 1). We found only three accessions (together < 1%) classified as CS, CR or CSRs. The accessions were mainly R-oriented: R, R/CR, R/CSR and R/SR represented 84% and 91% of all accessions in PHENOPSIS and Greenhouse, respectively (Table 1). Although we calculated CSR scores with only three leaf traits using the Pierce’s method, whole-plant traits were consistent with our classification. For instance, the C-axis is expected to be related to plant size and height, while the R-axis is expected to be related to flowering time and seed dispersal (Grime 1977; Hodgson *et al.* 1999). Accordingly, we found that C and R-axis were positively but poorly correlated with rosette fresh mass and the total number of fruits, respectively (ρ < 0.50, *P* < 0.05; Fig. S3).

**Table 1.** Proportion (%) of ecological strategies among *A. thaliana* accessions.

|  | *PHENOPSIS* | *Greenhouse* | *Original scores*<br>*from May et al.* | *Recalculated*<br>*scores with data*<br>*from May et al.*<br>*2017* |
| --- | --- | --- | --- | --- |
| **R** | 25.5 | 24.2 |  | 31.3 |
| **R/CR** | 47.6 | 58.6 | 12.5 |  |
| **R/CSR** | 8.7 | 7.1 |  |  |
| **R/SR** | 2.2 | 1.5 | 6.2 | 6.2 |
| **SR/CSR** | 8.4 | 4.5 |  | 12.5 |
| **SR** | 0.6 | 3.5 | 56.3 | 25.0 |
| **S** | 0.2 |  |  |  |
| **S/CSR** | 3.1 | 0.6 |  | 18.7 |
| **S/SC** |  |  | 25.0 |  |
| **S/SR** | 2.8 |  |  | 6.3 |
| **CSR** | 0.3 |  |  |  |
| **CS** | 0.3 |  |  |  |
| **CR** | 0.3 |  |  |  |

**Figure 1.**
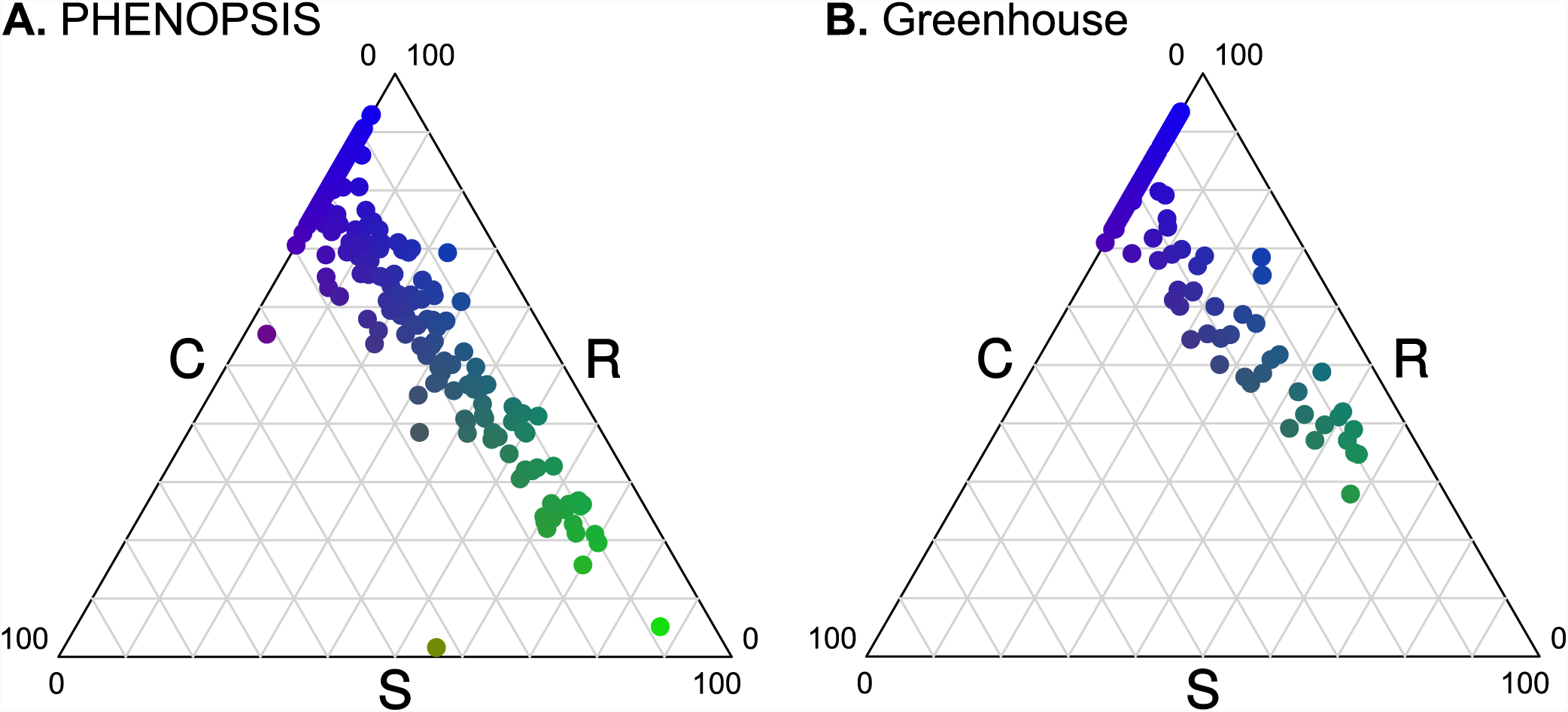
CSR variation in *A. thaliana*. (**A**) CSR representation of the 357 accessions from PHENOPSIS. (**B**) CSR representation of the 198 accessions from Greenhouse. Dots are coloured by CSR scores, following colour code provided in Pierce *et al.* (2017).

CSR scores were significantly correlated between PHENOPSIS and Greenhouse experiments, as measured across the 152 accessions common to both experiments (ρ = 0.34, 0.41 and 0.54 for C, S and R, respectively, all *P* < 0.001; Fig. S4). However, they were also significantly different between the two experiments (*P* < 0.01 for all the three scores). Accordingly, 78 accessions (51%) were classified in different CSR groups between the two experiments (“plastic” accessions hereafter). Globally, plastic accessions shifted towards more ruderal strategies in Greenhouse compared to PHENOPSIS, as reflected by the differences of S and R score between experiments (Fig. 2). 22% of the plastic accessions were classified as R in PHENOPSIS and R/CR in Greenhouse (inversely, 18% were classified as R/CR in PHENOPSIS and R in Greenhouse). Comparatively, C scores did not differ a lot between the two experiments (Fig. 2B).

**Figure 2.**
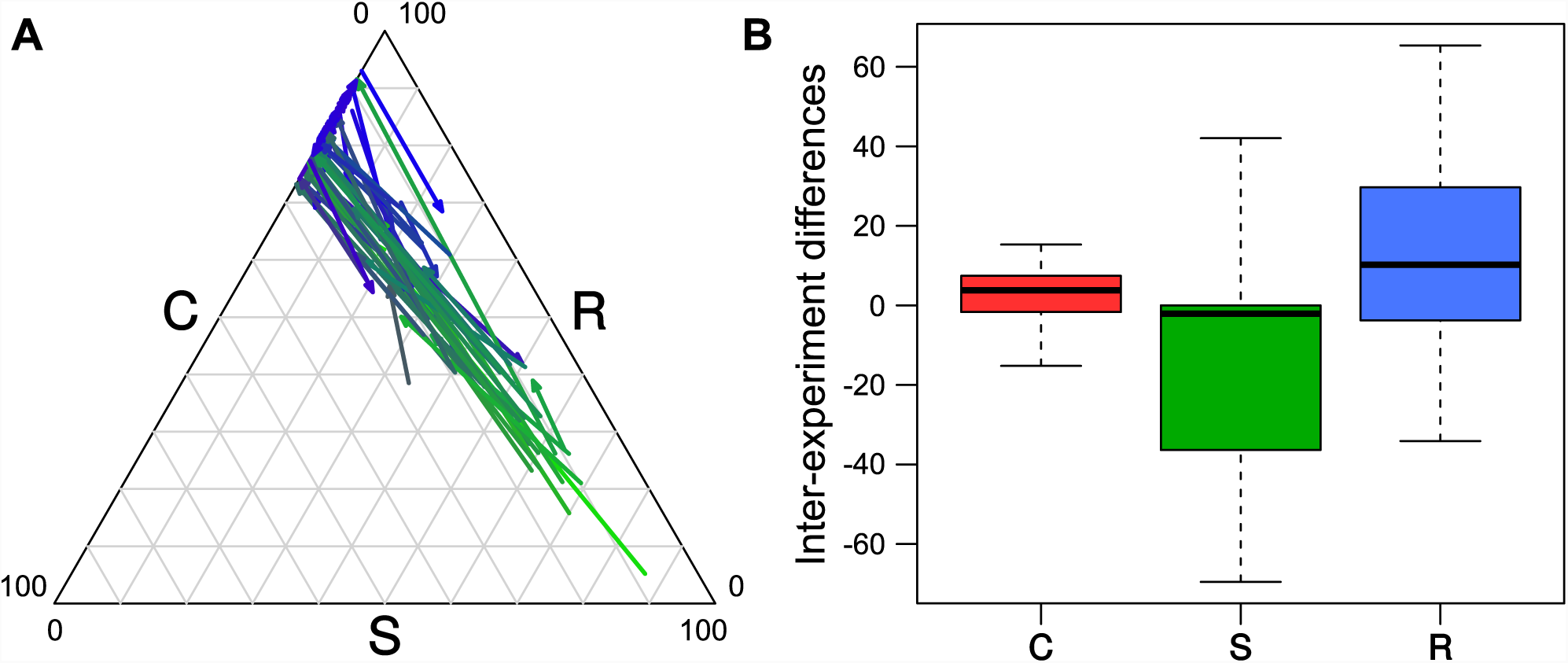
Plasticity of CSR classification in *A. thaliana*. (**A**) The 78 plastic accessions that have different CSR classification between PHENOPSIS and Greenhouse experiments are plotted. Arrows start at Greenhouse position and end at PHENOPSIS position, they are coloured according to CSR scores in PHENOPSIS, following colour code provided in Pierce *et al.* (2017). (**B**) Boxplot representing the difference in CSR scores between experiments (Greenhouse values – PHENOPSIS values).

### Relationships between CSR scores, flowering time and climate

Ruderality was positively correlated with SLA and mean annual temperature (MAT, °C) at the collection point of the accessions, but negatively with FT and LDMC (Fig. 3; Tables S4,S5). Thus, our results suggest that ruderality is typical of early-flowering plants with leaf traits representative of fast resource acquisition as reflected by low LDMC and high SLA values (Wright *et al.* 2004; Shipley *et al.* 2006). Inversely, stress-tolerators were characterized by late-flowering, with resource-conservative trait values such as high LDMC and low SLA; which were negatively correlated with MAT (Fig. S5). Consistently, S and R strategies were positively and negatively correlated with latitude, respectively (Table S4).

**Figure 3.**
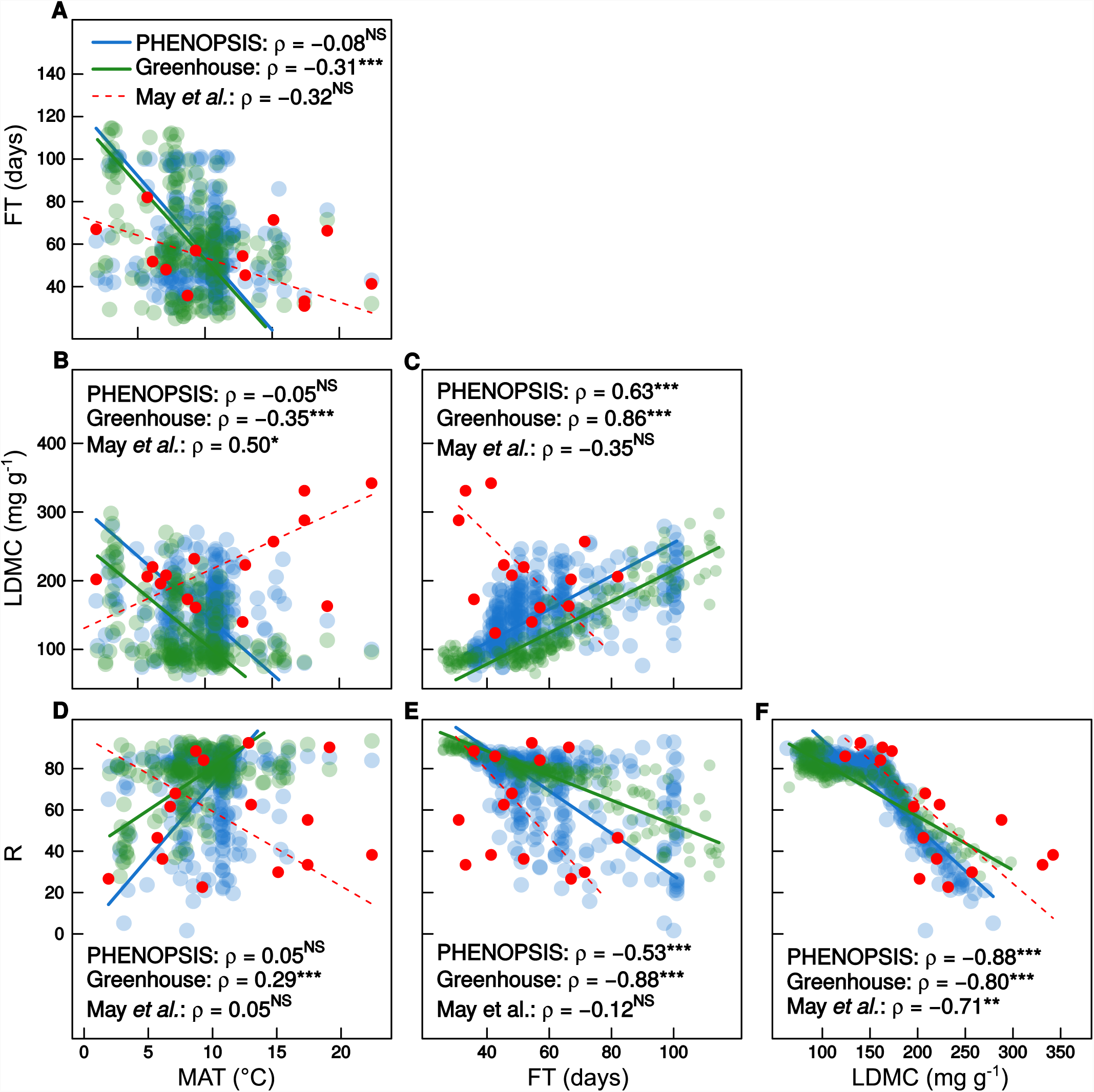
Relationships between ruderality, traits and environment in *A. thaliana*. Leaf trait and flowering time data were obtained from May *et al*. (2017) (red dots, *n* = 16), PHENOPSIS (green dots, *n* = 357) and Greenhouse (blue dots, *n* = 198). Ruderality (R) was calculated with Pierce’s method (2017) for all data. Mean annual temperature (MAT, °C) was extracted at the collection point of the accessions with Worldclim. FT: flowering time (days), LDMC: leaf dry matter content (mg g^−1^). ρ represents Spearman’s coefficient of correlation. Significance code: ^NS^: *P* > 0.1, : *P* < 0.1, *: *P* < 0.05, **: *P* < 0.01,***; *P* < 0.001. Regression lines were drawn from standard major axis (SMA).

Q_st_-F_st_ analysis suggested that the latitudinal variations of CSR strategies resulted from adaptive processes such as natural selection acting on leaf traits. Indeed, a value of Q_st_ significantly higher than F_st_ at neutral loci is generally considered as a signature of diversifying selection on the underlying traits (Leinonen *et al.* 2013). Here, we used the 95^th^ quantile of the F_st_ distribution genome-wide as a threshold of significance for phenotypic differentiation above neutral expectation. In Greenhouse, both S and R scores were significantly above neutral F_st_ (Q_st_ = 0.95, 95% CI = [0.72-1.00], and Q_st_ = 0.82, 95% CI = [0.62-1.00] for S and R, respectively, while mean F_st_ = 0.09 and F_st_ 95^th^ quantile = 0.35; Fig. 4A). In PHENOPSIS, only R scores were above, but non-significantly, neutral F_st_ (Q_st_ = 0.37 versus Fst 95^th^ quantile = 0.33). S scores were slightly, and non-significantly, below the neutral expectation (Q_st_ = 0.29, 95% CI = [0.10-0.80]; Fig. 4C). By contrast, in both Greenhouse and PHENOPSIS, Q_st_ of C scores were close to 0, suggesting that this axis of plant strategies did not vary under the influence of adaptive processes in *A. thaliana*. The lower Q_st_ values reported for the PHENOPSIS experiment can be explained by the absence of individual replicates in this experiment. By contrast, using genotypic mean in Greenhouse across four replicates allowed reducing intra-genotypic variance, and thus total phenotypic variance compared to phenotypic variance between genetic groups. Consistent with these results, plotting the distribution of *A. thaliana* ecological strategies across Europe (Fig. 4B,D) revealed that accessions with S-oriented strategies (S, SR, SR/CSR, S/CSR, S/SC, SC and SC/CSR) were originating from northern regions, Sweden in particular.

**Figure 4.**
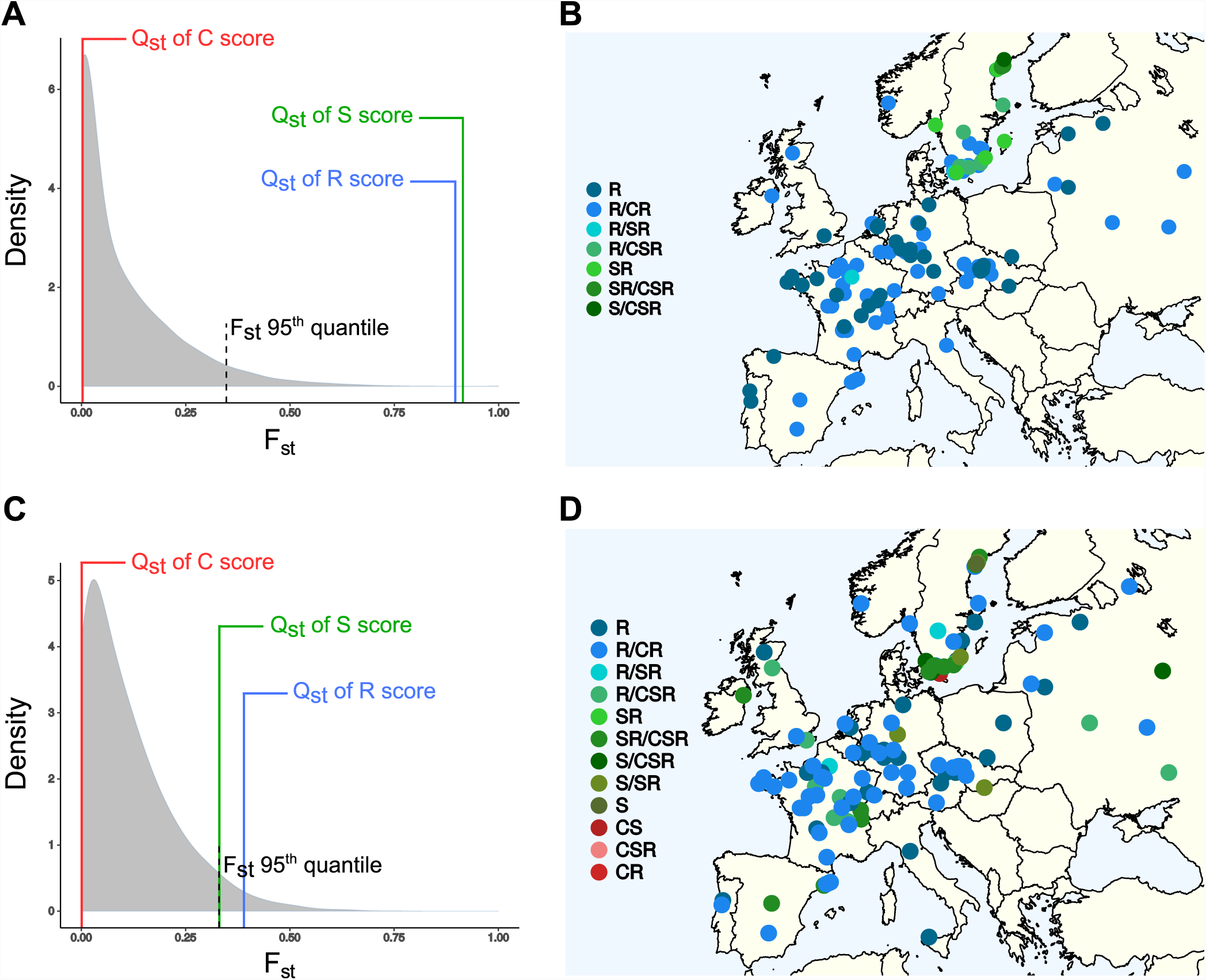
Q_st_-F_st_ analysis and geographic location of CSR strategies in *A. thaliana*. Distribution (in grey) of F_st_ values across the 24,562 SNPs with 95^th^ quantile threshold of non-neutral expectation (dashed line), and Q_st_ values for the C, S and R scores measured as the ratio of phenotypic variance between genetic groups over total phenotypic variance. Analysis performed independently on Greenhouse dataset (**A**), and PHENOPSIS dataset (**C**). Geographic location of CSR strategies with Greenhouse dataset (**B**, *n* = 198), and PHENOPSIS dataset (**D**, *n* = 357).

### Comparison with observations from May *et al*

In contrast with our results, the 16 accessions in the study published by May *et al.* were mainly categorized as S-oriented: S/SC, S/SR, S/CSR, SR and SR/CSR (Table 1, Fig. S6). May *et al.* used the Hodgson’s method to calculate CSR scores with seven traits, including flowering time and duration, two important components of ruderality (Hodgson *et al.* 1999). To compare the two classification methods, we used the traits values for LA, SLA and LDMC provided in May *et al*. (2017) to calculate CSR scores with the Pierce’s method and compared them with the Hodgson’s method. The CSR scores calculated with the two methods were positively correlated (ρ = 0.77, 0.79 and 0.73 for C, S and R, respectively; all *P* < 0.01; Fig. S7), suggesting that both methods return similar categorization (Table 1). However and despite the significant correlations, CSR scores varied substantially between the two methods (Fig. S7). This showed that the traits related to ruderality (flowering time and duration) and competition (plant height and lateral spread) used in the Hodgson’s method impacted the inference of plant ecological strategies compared to leaf traits alone.

FT measured in this study was strongly positively correlated with FT measured in May *et al.* under controlled conditions (*n* = 10 and 6 in PHENOPSIS and Greenhouse, respectively, both *r* = 0.96, *P* < 0.01; Fig. S8A). By contrast, LDMC measured in May *et al.* was negatively correlated with our measurements (Fig. S8D), possibly because of three individuals with early FT and extremely high LDMC values (> 250 mg g^−1^) compared to our measurement (< 110 mg g^−1^) on the same accessions (red dots in Fig. 3C). As a result, FT and LDMC were negatively, albeit non-significantly, correlated in the May *et al.* study (ρ = −0.35; Fig. 3C). Furthermore, there was a positive correlation between LDMC and MAT in May *et al.* (*P* < 0.05), while we found the opposite in both PHENOPSIS and Greenhouse (Fig. 3B). By construction of the CSR classification method, LDMC strongly participates to S and R axes (Fig. 3F and Fig. S5). Consequently, the positive correlation between LDMC and MAT found in May *et al.* was associated with a positive correlation between S and MAT (Fig. S5D), and inversely a negative correlation between R and MAT (Fig. 3D), although these two relationships were not significant with the 16 accessions from May *et al.* when using the Pierce’s method of CSR classification.

## Discussion

### Functional adaptation to climate in *A. thaliana*

The relationship between CSR and climate at the interspecific level is still not well established (Pierce *et al.* 2017). More broadly, trait-environment relationships remain a central question in functional ecology and functional biogeography (Poorter *et al.* 2009; Violle *et al.* 2014; Borgy *et al.* 2017; Butler *et al.* 2017; Šímová *et al.* 2018). By contrast, adaptation to climate has been widely studied within species, notably genetic adaptation along latitudinal or altitudinal gradients in annual plants, and in *A. thaliana* in particular (Johanson *et al.* 2000; Picó *et al.* 2008; Banta *et al.* 2012; Guo *et al.* 2012; Brachi *et al.* 2013; Wolfe and Tonsor 2014; Bloomer and Dean 2017; Tabas-Madrid *et al.* 2018). Indeed, *A. thaliana* has been the model species in molecular biology, plant genetics and evolution for the last decades (Bergelson and Roux 2010; Weigel 2012). It is widely distributed in various climates, but is generally considered as a ruderal species that grows fast, reproduces early and dies right after seed dispersal (Pierce *et al.* 2017). As expected, we found in this study that *A. thaliana* was predominantly ruderal, secondly stress-tolerator and poorly competitor. However, we showed an important range of CSR variation among *A. thaliana* accessions along the S-R axis and associated with flowering time variation.

Consistent with previous studies, flowering time was positively correlated with latitude (Caicedo *et al.* 2004; Lempe *et al.* 2005; Banta *et al.* 2012). For instance, northern accessions exhibit late-flowering and long life cycle even when they are grown under controlled conditions in growth chamber or greenhouse (Vasseur *et al.* 2018). Our results showed that flowering time was positively correlated with LDMC, and that values for the two traits were higher in accessions originating from higher latitude and lower temperatures. Thus, northern accessions exhibit a suite of traits associated with resource conservation and longevity such as late flowering, high LDMC and low SLA (Wright *et al.* 2004; Shipley *et al.* 2006; Vasseur *et al.* 2012). Q_st_-F_st_ analysis revealed that these latitudinal variations result from the adaptive diversification of leaf traits. These adaptive shifts can be explained because, in cold regions, biomass production during the growing season is limited by various stresses. Low temperatures directly limit plant growth rate by slowing down metabolic processes. Furthermore, cold indirectly limits plant growth rate because of the reduction in the availability of water and nutrients. In these conditions, slow growing genotypes with long life cycle, associated with high LDMC, low SLA and low metabolic activities, can be an efficient strategy. Interestingly, stress tolerance has been shown to be selected at both ends of the geographic range of *A. thaliana*, but is expressed under different temperature conditions (Exposito-Alonso *et al.* 2018; Vasseur *et al.* 2018).

Conversely, ruderal strategies were more abundant in temperate and hot environments. Ruderal plants are typically associated with a short life cycle, low LDMC and high SLA, and presumably high metabolic rate and low tissues protection (Grime 1977). In temperate climates with a relatively long growing season and high resource availability, these characteristics may allow *A. thaliana* individuals to complete their growth cycle early and avoid competition with taller species. Furthermore, in hot and dry climates with a shorter growing period (*e.g.*, Mediterranean climate), fast growing strategies may allow *A. thaliana* individuals to complete their growth cycle and disperse before the onset of drought, which operate as a disturbance rather than a stress, and should therefore be more favourable to ruderality (Madon & Médail 1997; Volaire 2018). This result is consistent with interspecific studies at global scale that reported a positive relationship between SLA and temperature in herbaceous species (Borgy *et al.* 2017; Šímová *et al.* 2018). This can be interpreted as a sign of selection for fast-growth, ruderal strategies in hot and stressing environments at both intra- and interspecific levels (Anderegg *et al.* 2018).

The lack of adaptive differentiation between genetic groups along the C axis, as reflected by the low Q_st_ values compared to neutral F_st_, can be explained by the low variation of competitive ability among *A. thaliana* accessions. Additionally, it could suggest that competitive environments can be found in various climates as long as stress does not dominate vegetation processes. This would also explain the lack of a clear geographic pattern and latitudinal gradient of competitive ability across plant populations and species (Damgaard and Weiner 2017).

### Influence of classification methodologies, trait measurement and growth conditions on trait-environment relationships

Trait-trait, trait-CSR and trait-environment relationships were sometimes opposite between May *et al.* (2017) and our study. For instance, May *et al.* (2017) reported a positive correlation between stress-tolerance and mean temperature, while we found the opposite. A first explanation of these differences is the methods used to calculate CSR scores among accessions. Although Pierce’s and Hodgson’s scores were all positively correlated when performed on the same set of traits and accessions, scores obtained from the two methods varied substantially. For instance, an accession had S score at 35% with Hodgson’s method but 0% with Pierce’s method (Fig. S7B). The re-analysis of May *et al.* data made by the authors (A. Wingler, personal communication) indicated that the three accessions with very high values for S identified using the Hodgson’s method (Mt-0, Cvi-0 and Ct-1) were no longer in the top three ranked accessions for S when using the Pierce’s method, which led to a lack of correlation of S and R with temperature when using this method. This can be explained because life history traits at whole-plant level, notably flowering time and plant size, are important components of ruderality and competitive ability in herbaceous species (Violle *et al*. 2009; Hodgson *et al*. 2017), but they are not included in the Pierce’ method of CSR classification. Here, we found that C and R axes calculated with leaf traits were positively, but poorly, correlated with rosette fresh mass and fruit number, respectively. Additionally, many early-flowering accessions were similarly classified as purely ruderal (R = 100%,), although they displayed variations in leaf traits and flowering time, and consequently, in their level of ruderality. This was translated into no, or small differences in CSR strategies between accessions from temperate and Mediterranean climates (Fig. 4), although Mediterranean accessions can be very short-lived and thus, more ruderal than accessions from less stressing environments (Vasseur *et al*. 2018). Together, this suggests that classification methods based on leaf traits can be powerful to screen large database or to perform many measurements at global scale, but it might be limited to examine subtle variations within species and/or in specific taxa. For instance, including other, easily-measurable traits might be necessary to better describe ruderality in annual plants, such as phytomer miniaturization and the number of juvenile phytomers, because each promotes early maturity (Hodgson *et al.*, 2017).

A second explanation to the opposite trait-environment relationships found between this study and May *et al*. is the difference in the protocols used for trait measurement. In our experiments, we followed the recommended procedures to phenotype traits of all individuals at the same *ontogenetic stage* (Reich *et al.* 1999; Perez-Harguindeguy *et al.* 2013). Specifically, LDMC and SLA were measured at the transition to flowering (*i.e.* bolting stage). By contrast, leaf traits were measured in a growth chamber at the same *age* by May *et al.* (61 days for LDMC), although flowering time in growth chamber varied from 30 days to 82 days (and some accessions did not flower at all), and although it is widely recognized that leaf traits strongly vary during plant ontogeny (Walters *et al.* 1993; Hérault *et al.* 2011; Pantin *et al.* 2012). In other words, LDMC was measured 30 days after flowering for the earliest accessions and before flowering for the latest ones. With such a procedure, the leaves compared might have been in contrasted physiological stages. In particular, leaves measured on the early-flowering accessions might have been - at least in part - senescing, which may result in much higher LDMC values - and lower SLA values - in these accessions (Fig. 3C). In agreement with this hypothesis, the LDMC values measured on the early-flowering accessions in our experiment were approximately half of the values estimated by May *et al.*. As LDMC strongly participates to the S-R axis, this could explain the opposite correlations between CSR and environment between the two studies. Furthermore, we found that FT was positively correlated with LDMC, consistently with previous studies in a smaller set of accessions (Vile *et al.* 2012), as well as in recombinant inbred lines (El-Lithy *et al.* 2010; Vasseur *et al.* 2012, 2014). Previous studies have notably reported that early-flowering genotypes have resource-acquisitive strategies, characterized by high SLA but low LDMC and short lifespan (El-Lithy *et al.* 2010; Vasseur *et al.* 2012, 2014, 2018; Blonder *et al.* 2015).

Finally, opposite correlations between studies might also partly result from trait plasticity to growth conditions. FT in *A. thaliana* is expected to vary with light conditions and temperature (Mouradov *et al.* 2002). For instance, *A. thaliana* does not generally flower under short-day conditions. In our study, traits were measured in controlled and constant conditions, on plants grown in 12 h photoperiod and without cold exposure (*i.e.* vernalization). However, we could expect FT and leaf traits, and thus CSR-environment relationships, to be different when measured on plants grown outside like in May *et al.*, after vernalization or in short-or long-day conditions. Consistent with this idea, we found that half the accessions common to PHENOPSIS and Greenhouse did not have the same position in the CSR space: plants grown in the greenhouse were generally shifted towards the R end of the spectrum compared to plants grown in PHENOPSIS. This can be explained by the relative low light intensity provided by artificial lamps in the greenhouse compared to the phenotyping platform (65 versus 175 μmol m^−2^ s^−1^ PPFD). In addition, plants were grown in the greenhouse at higher density than in PHENOPSIS, which could have increased competition for light between plants. The shade-avoidance syndrome has been described as a suite of leaf trait responses to low light and competition (Kim *et al.* 2005; Mullen *et al.* 2006). This includes an increase in leaf angle and SLA, associated with a reduction in LDMC and flowering time (Kim *et al.* 2005; Vasseur *et al.* 2011). This is consistent with a shift towards resource-acquisitive strategies in Greenhouse. Importantly and more broadly, controlled conditions are very different from natural conditions that plants experience in the wild, and where plants should ideally be measured to properly infer their ecological strategies. However, it remains difficult to take into account genotype-by-environment interactions when screening genotypes in natural conditions. Consequently, trait-based approaches for functional classification of plants were initially proposed as a tool to infer the adaptive significance of traits in controlled conditions (Grime and Hunt, 1975).

## Conclusion

Intraspecific variation in functional strategies varied substantially along the S-R axis in *A. thaliana*. Tolerance to stress seems to be favoured in cold environments at higher latitudes while ruderality is predominant in temperate and hot climates. However, CSR categorization within species, specifically in an herbaceous species like *A. thaliana*, is sensitive to several parameters such as the type of traits used to classify accessions and the protocols used for trait measurement. Furthermore, our results suggest that phenotypic plasticity to growth conditions can significantly impacts trait values and thus, the determination of plant ecological strategies. This suggests that the use of trait databases for local or global analyses of trait-environment relationships at species level might suffer from biases due to both phenotypic plasticity and intraspecific trait variation. In a recent analysis, ruderality has been demonstrated to correlate positively with the probability of naturalization of alien species (Guo *et al.* 2018). In future studies, it will be interesting to examine in more details the response of traits, trait combinations and strategies to environmental conditions. For instance, analysing the plasticity of CSR strategies to different temperatures and water stresses could reveal whether S-related strategies are constitutive or stress induced, and whether invasive species show greater plasticity in ecological strategies than other species.

## Acknowledgments

We thank Myriam Dauzat for her technical assistance during trait measurement and environmental control of the PHENOPSIS platform. We also warmly thank Astrid Wingler and John Hodgson for their helpful comments and suggestions as referees of a previous version of the present manuscript. EB, KS, CV and DV performed the experiments, FV, DV and CV performed statistical analyses. All authors interpreted the results. FV wrote the manuscript with help from all authors.

**List of Supplementary Information**

**Table S1. Phenotypic traits measured in PHENOPSIS experiment.**

**Table S2. Phenotypic traits measured in Greenhouse experiment.**

**Table S3. Heritability and genetic effects on traits measured in Greenhouse experiment.**

**Table S4. Spearman’s pairwise correlations between traits and environments.**

**Table S5. Pearson’s pairwise correlations between traits and environments.**

**Figure S1. Geographic location of the genetic groups defined with SNPs clustering.** (**A**) Representation of the six genetic groups (+ admixed) identified in the PHENOPSIS dataset (*n* = 394). (**B**) Representation of the four genetic groups (+ admixed) identified in the Greenhouse dataset (*n* = 198).

**Figure S2. Trait-trait relationships in *A. thaliana*.** PHENOPSIS (blue dots, *n* = 357) and Greenhouse (green dots, *n* = 198). ρ represents Spearman’s coefficient of correlation. Significance code: ^NS^: *P* > 0.1, : *P* < 0.1, *: *P* < 0.05, **: *P* < 0.01,***; *P* < 0.001. Regression lines were drawn from standard major axis (SMA).

**Figure S3. Correlations between C and R axes, plant biomass and fruit number.** (**A**) Correlation between C axis and rosette fresh weight (mg) measured in PHENOPSIS (blue dots, *n* = 357). (**B**) Correlation between R axis and fruit number measured in Greenhouse (green dots, *n* = 198). ρ represents Spearman’s coefficient of correlation. Significance code: ^NS^: *P* > 0.1, : *P* < 0.1, *: *P* < 0.05, **: *P* < 0.01,***; *P* < 0.001. Regression lines were drawn from standard major axis (SMA).

**Figure S4. Correlations between CSR scores in PHENOPSIS and Greenhouse experiments.** Correlations were estimated on the set of 152 accessions common to both experiments. ρ represents Spearman’s coefficient of correlation. Significance code: ^NS^: *P* > 0.1, : *P* < 0.1, *: *P* < 0.05, **: *P* < 0.01,***; *P* < 0.001. Regression lines were drawn from standard major axis (SMA).

**Figure S5. CSR-traits and CSR-environment relationships in *A. thaliana*.** Leaf trait and flowering time data were obtained from May *et al*. (2017) (red dots, *n* = 16), PHENOPSIS (blue dots, *n* = 357) and Greenhouse (green dots, *n* = 198). Ruderality (R), stress-tolerance (S), and competitive ability (C) were calculated with Pierce’s method (2017) for all data. Mean annual temperature (MAT, °C) was extracted at the collection point of the accessions with Worldclim. FT: flowering time (days), LDMC: leaf dry matter content (mg g^−1^). ρ represents Spearman’s coefficient of correlation. Significance code: ^NS^: *P* > 0.1, : *P* < 0.1, *: *P* < 0.05, **: *P* < 0.01,***; *P* < 0.001. Regression lines were drawn from standard major axis (SMA).

**Figure S6. CSR representation of the 16 accessions from May *et al.* (2017).** CSR scores recalculated with leaf trait data using Pierce’s (2017) method. Dots are coloured by CSR scores, following colour code provided in Pierce *et al.* (2017).

**Figure S7. Correlation between Hodgson’s and Pierce’s methods for quantifying CSR.** CSR scores were recalculated with leaf traits provided for the 16 accessions in May *et al.* (2017), using Pierce’s method, and compared to CSR scores measured in May *et al.* (2017) with Hodgson’s method. ρ represents Spearman’s coefficient of correlation. Significance code: ^NS^: *P* > 0.1, : *P* < 0.1, *: *P* < 0.05, **: *P* < 0.01,***; *P* < 0.001. Regression lines were drawn from standard major axis (SMA).

**Figure S8. Correlations between traits measured in May *et al.* and the present study.** Correlations were estimated on the 10 and 6 accessions common to May *et al*. (2017) and PHENOPSIS (blue dots) and Greenhouse (green dots) experiments, respectively. ρ represents Spearman’s coefficient of correlation. Significance code: ^NS^: *P* > 0.1, : *P* < 0.1, *: *P* < 0.05, **: *P* < 0.01,***; *P* < 0.001. Regression lines were drawn from standard major axis (SMA). FT: flowering time (days), LA: leaf area (mm^2^), LDMC: leaf dry matter content (mg g^−1^), SLA: specific leaf area (m^2^ kg^−1^).

